# Activation of endogenous retroviruses during brain development causes neuroinflammation

**DOI:** 10.1101/2020.07.07.191668

**Authors:** Marie E Jönsson, Raquel Garza, Yogita Sharma, Rebecca Petri, Erik Södersten, Jenny G Johansson, Pia A Johansson, Diahann AM Atacho, Karolina Pircs, Sofia Madsen, David Yudovich, Ramprasad Ramakrishnan, Johan Holmberg, Jonas Larsson, Patric Jern, Johan Jakobsson

**Affiliations:** Laboratory of Molecular Neurogenetics, Department of Experimental Medical Science, Wallenberg Neuroscience Center and Lund Stem Cell Center, BMC A11, Lund University, 221 84 Lund, Sweden; Department of Cell and Molecular Biology, Karolinska Institutet, Solnavägen 9, 171 65, Stockholm, Sweden; Division of Molecular Medicine and Gene Therapy, Department of Laboratory Medicine and Lund Stem Cell Center, BMC A12, Lund University, 221 84, Lund, Sweden; Division of Clinical Genetics, Lund University, 22184, Lund, Sweden; Science for Life Laboratory, Department for Medical Biochemistry and Microbiology, Uppsala University, 751 23 Uppsala, Sweden

## Abstract

Endogenous retroviruses (ERVs) make up a large fraction of mammalian genomes and are thought to contribute to human disease, including brain disorders. In the brain, aberrant activation of ERVs is a potential trigger for neuroinflammation, but mechanistic insight into this phenomenon remains lacking. Using CRISPR/Cas9-based gene disruption of the epigenetic co-repressor protein Trim28, we found a dynamic H3K9me3-dependent regulation of ERVs in proliferating neural progenitor cells (NPCs), but not in adult neurons. *In vivo* deletion of *Trim28* in cortical NPCs during mouse brain development resulted in viable offspring expressing high levels of ERV expression in excitatory neurons in the adult brain. Neuronal ERV expression was linked to inflammation, including activated microglia, and aggregates of ERV-derived proteins. This study demonstrates that brain development is a critical period for the silencing of ERVs and provides causal *in vivo* evidence demonstrating that transcriptional activation of ERV in neurons results in neuroinflammation.

## Introduction

About one tenth of the human and mouse genomes is made up of endogenous retroviruses (ERVs) (Jern and Coffin, 2008). This is a result of the cumulative infection of the germ line by retroviruses over millions of years. ERVs are dynamically silenced at the transcriptional level during early development via epigenetic modifications, including histone methylation and deacetylation as well as DNA methylation (Rowe et al., 2010; Yoder et al., 1997). Together, these repressive mechanisms suppress ERV expression in somatic tissues. However, it is becoming increasingly clear that ERVs are aberrantly activated in various human diseases, including a number of neurological disorders. For example, ERV expression has been found to be elevated in the cerebrospinal fluid and in post-mortem brain biopsies from patients with multiple sclerosis, amyotrophic lateral sclerosis, Alzheimer’s disease, Parkinson’s disease and schizophrenia (Andrews et al., 2000; Douville et al., 2011; Garson et al., 1998; Guo et al., 2018; Karlsson et al., 2001; Li et al., 2015; MacGowan et al., 2007; Perron et al., 1997; Perron et al., 2008; Steele et al., 2005; Sun et al., 2018; Tam et al., 2019b).

Aberrant activation of ERVs in the brain has been proposed to be directly involved in the disease process through a number of different mechanisms, including the activation of an innate immune response, direct or indirect neurotoxicity or by modulating endogenous gene networks (Jonsson et al., 2020; Saleh et al., 2019; Tam et al., 2019a). However, causal studies of ERV activation in the brain are challenging, since this phenomenon is difficult to model in the lab. Most experimental studies rely on ectopic expression of ERV-derived transcripts, often using xeno-overexpression at non-physiological levels, making it hard to interpret the results (see e.g. (Antony et al., 2004; Li et al., 2015)). Still, while the role of ERVs in neurological disorders remains unclear (Tam et al., 2019a), ERV-activation may constitute a new type of disease mechanism that could be exploited to develop much needed therapy for these disorders. Direct experimental evidence on the mechanisms underlying ERV repression and the consequences of ERV activation in the brain is therefore required.

We recently found that Trim28, an epigenetic co-repressor protein, silences ERV expression in mouse and human neural progenitor cells (NPCs) (Brattas et al., 2017; Fasching et al., 2015). Trim28 is recruited to genomic ERVs via Krüppel-associated box - zinc finger proteins (KRAB-ZFPs), a large family of sequence-specific transcription factors (Imbeault et al., 2017). Trim28 attracts a multiprotein complex that establishes transcriptional silencing and deposition of the repressive histone mark H3K9me3 (Sripathy et al., 2006). Trim28 is highly expressed in the brain and has been linked to behavioral phenotypes reminiscent of psychiatric disorders (Fasching et al., 2015; Jakobsson et al., 2008; Whitelaw et al., 2010).

In this study, we have investigated the consequences of *Trim28* deletion in the developing and adult mouse brain. We found that while Trim28 is needed for the repression of ERVs during brain development it is redundant in the adult brain. Our results demonstrate the presence of an epigenetic switch during brain development, where the dynamic Trim28-mediated ERV repression is replaced by a different more stable mechanism. Interestingly, conditional deletion of *Trim28* during brain development resulted in ERV expression in adult neurons leading to neuroinflammation, including the presence of activated microglia and aggregates of ERV-derived proteins. In summary, our results provide direct experimental evidence *in vivo* for a link between aberrant ERV expression in the brain and the presence of neuroinflammation.

## Results

### CRISPR/Cas9-mediated deletion of Trim28 in mouse NPCs

To evaluate the consequences of acute loss of *Trim28* in NPCs, we used CRISPR-Cas9 gene disruption. We generated NPC cultures from Rosa26-Cas9 knock-in transgenic mice (Fig 1a) (Platt et al., 2014), in which Cas9-GFP is constitutively expressed in all cells. These Cas9-NPCs were transduced with a lentiviral vector expressing gRNAs (LV.gRNAs) designed to target either exon 3, 4 or 13 of *Trim28* (g3, g4, g13) or to target *lacZ* (control). The vector also expressed a nuclear RFP reporter gene (H2B-RFP). Cas9-NPCs transduced with LV.gRNAs were expanded for 10 days, at which point RFP expressing cells were isolated by FACS (Fig 1a). To assess gene editing efficiency, we extracted genomic DNA from the RFP+ Cas9-NPCs and performed DNA amplicon sequencing of the different gRNA target sites. We found that all three gRNAs (g3, g4 and g13) were highly effective, generating indels at a frequency of 98-99% at their respective target sequences (Fig 1b). The majority of these indels caused a frameshift in the Trim28 coding sequence that is predicative of loss-of-function alleles (Fig 1b). These results demonstrate that CRISPR/Cas9-mediated gene disruption is an efficient way to investigate the functional role of *Trim28*.

**Figure 1.**
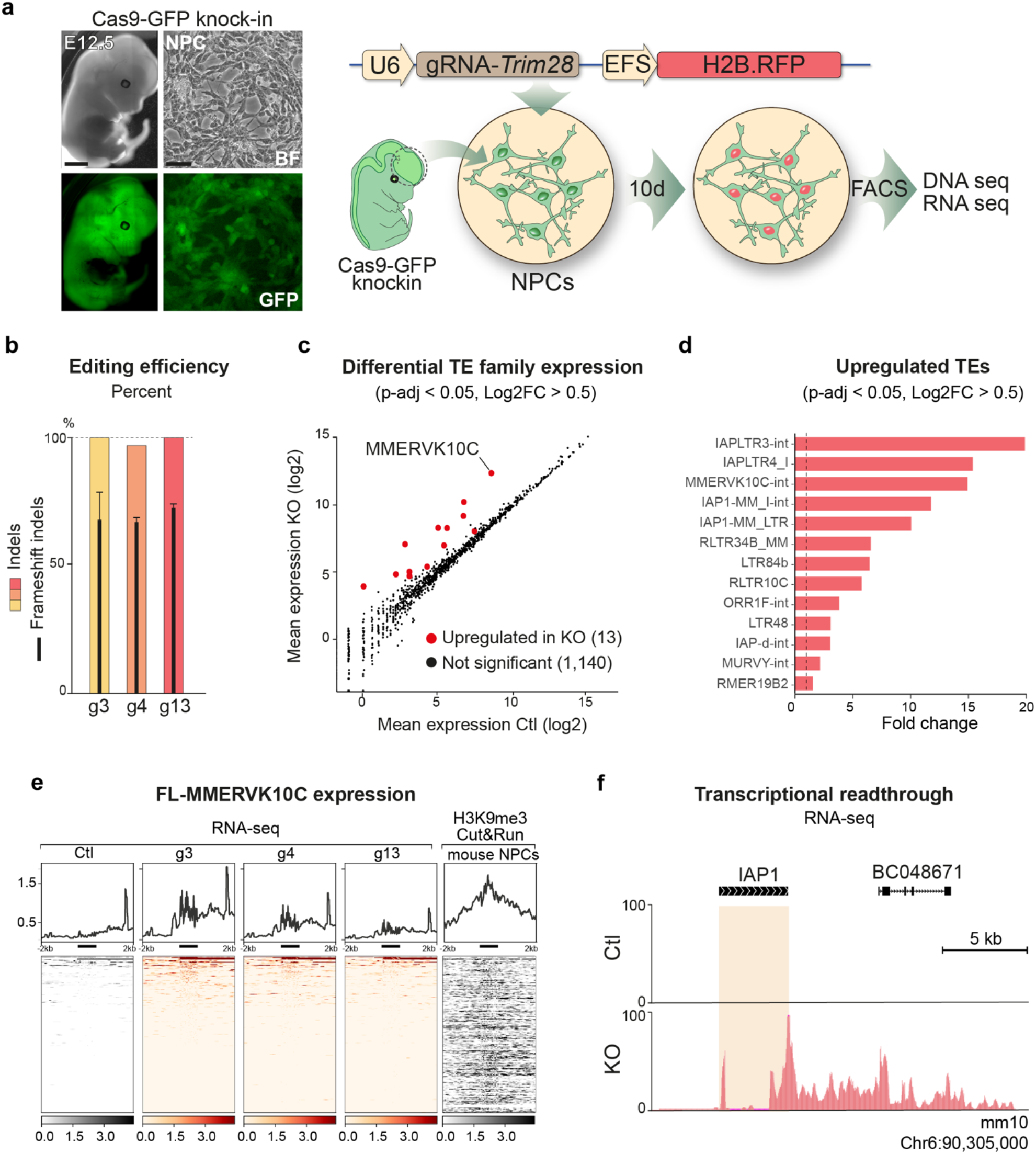
CRISPR/Cas9 based deletion of Trim28 in NPCs results in upregulation of ERVs. (a) A schematic of the workflow for Trim28-KO in mouse NPC cultures. Scalebars: embryo 1 mm, NPCs 20 μm. (b) Estimation of gene editing at the Trim28 loci using NGS-sequencing of amplicons. Black bars indicate % of frameshift indels (c) RNA-seq analysis of the expression of TE-families using TEtranscripts (d) The significantly upregulated TE families with a fold change larger than 0.5 upon Trim28-KO in mouse NPCs. The dashed line indicates significance. (e) CUT&RUN analysis of H3K9me3 in mouse NPCs of all full length MMERVK10Cs (right panel) RNA-seq analysis of the Trim28-KO and control samples, visualising full length MMERVK10C elements. The location of the full length MMERVK10Cs are indicated as a thick black line under each histogram. (f) Example of transcriptional readthrough outside the full length MMERVK10Cs into a nearby gene.

### Acute loss of *Trim28* in mouse NPCs results in upregulation of ERVs

We next queried if acute *Trim28* deletion in NPCs influences the expression of ERVs and other transposable elements (TEs). We performed strand-specific 2×150bp RNA-seq on LV.gRNA transduced Cas9-NPCs and investigated the change of expression in different ERV-families using a TE oriented read quantification software, TEtranscripts (Jin et al., 2015), while individual elements were analysed using a unique mapping approach. Both of these analyses revealed an upregulation of ERVs upon the CRISPR-mediated *Trim28*-KO in mouse NPCs (Fig 1c-d, Fig S1a). We found 13 upregulated ERV families, including IAPs and MMERVK10C. Both IAPs and MMERVK10C are recent additions to the mouse genome and these ERV-families include many full-length, transposition-competent elements with the potential to produce long transcripts and ERV-derived peptides. We also investigated the expression of other classes of TEs such as LINE-1s and SINEs but found no evidence of upregulation. Thus, acute deletion of *Trim28* in NPCs causes a specific upregulation of ERVs.

*Trim28* attracts a repression complex containing the histone methyltransferase SETDB1 that deposits H3K9me3 (Sripathy et al., 2006). CUT&RUN epigenomic analysis (Skene and Henikoff, 2017) of this histone modification in NPCs revealed that *Trim28*-controlled ERVs were covered by H3K9me3, directly linking the activation of ERVs to *Trim28* binding. For example, almost all full-length members of the MMERVK10C family were covered by H3K9me3 (Fig 1e). Notably, only a handful of these individual elements were transcriptionally activated after *Trim28*-KO. This suggests that *Trim28* binds to many full-length ERVs in NPCs but is only responsible for transcriptional silencing of a small subset of them. However, these elements are highly expressed upon *Trim28* deletion.

TEs have the potential to change the surrounding epigenetic landscape and consequently influence the expression of protein-coding genes in their vicinity (Brattas et al., 2017; Chuong et al., 2017; Fasching et al., 2015). Accordingly, we found, for example, that genes located in the close vicinity of an upregulated MMERVK10C element also displayed a significant upregulation (Fig S1b). In some instances, this was due to the activated ERV acting as an alternative promoter (Fig 1f). Since Trim28 has additional functions in the cell (Quenneville et al., 2011) (Ziv et al., 2006) (Bunch et al., 2014), we also investigated the expression of all protein coding genes in *Trim28*-KO NPCs. This analysis revealed that acute loss of *Trim28* in NPCs had only a modest effect on protein coding genes (Fig S1c). Indeed, PCA analysis using protein coding genes was unable to distinguish *Trim28*-KO cells from control NPCs. By contrast, a PCA analysis using ERVs clearly separates the two groups (Fig S1d-e). These results demonstrate that Trim28 represses the transcription of ERVs in NPCs but has a marginal direct effect on protein coding genes.

### Deletion of *Trim28* in adult neurons *in vivo*

Alongside our previous data (Brattas et al., 2017; Fasching et al., 2015), these results confirm that *Trim28* is critical to repress ERVs in NPCs. However, it remains unclear how relevant these findings are to the situation *in vivo*. Therefore, we next investigated the consequences of deleting *Trim28* in mature neurons of adult mice. We designed adeno associated viral vectors (AAV) vectors to drive expression of the *in vitro* verified gRNAs (AAV.gRNA) and used them separately or combined in the forebrain of Cas9-GFP knock-in mice. The AAV.gRNA vectors expressed H2B-RFP under the neuronal specific Synapsin promoter, allowing us to visualize and isolate transduced neurons (Fig 2a).

**Figure 2.**
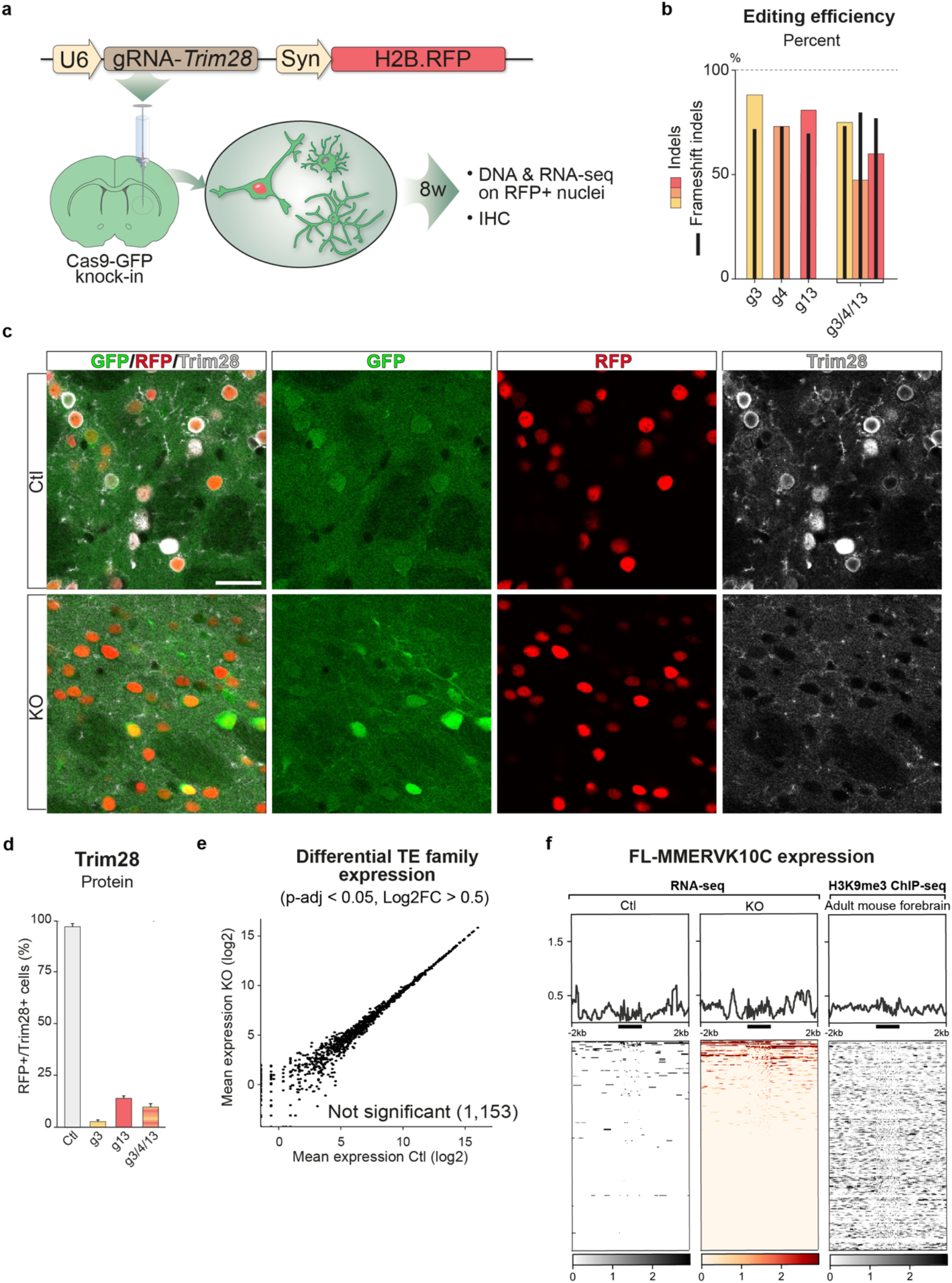
CRISPR-Cas9 deletion of Trim28 in adult neurons in vivo. (a) A schematic of the workflow targeting Trim28 in the mouse forebrain using AAV vectors expressing the gRNA and a nuclear RFP reporter. 8 weeks later the injected animals were analysed either by immunohistochemical analysis or nuclei isolation by FACS prior to DNA/RNA sequencing. (b) Estimation of gene editing at the Trim28 loci using NGS-sequencing of amplicons. Black bars indicate % of the detected indels that disrupted the frameshift. Scalebar 30 μm. (c-d) Gene editing of the Trim28-loci resulted in a robust loss of Trim28 protein, as evaluated by IHC where the expression of Trim28 in RFP+ cells were quantified. (e) RNA-seq analysis of the expression of TE-families using TEtranscripts. (f) ChIP-seq analysis of H3K9me3 in adult forebrain neurons of all full length MMERVK10Cs (right panel) RNA-seq analysis of the Trim28-KO and control samples, visualising full length MMERVK10C elements. The location of the full length MMERVK10Cs are indicated as a thick black line under each histogram.

We injected AAV.gRNA into the forebrain of adult Cas9-GFP mice and sacrificed the animals after eight weeks. The RFP expressing neuronal nuclei were isolated by FACS and amplicon sequenced to estimate the gene editing efficiency. All three gRNAs resulted in highly efficient gene editing (indel frequencies of 73-89%) where the majority of the indels were frameshift mutations (Fig. 2b). Efficient deletion of *Trim28* was subsequently verified by immunohistochemistry (IHC) analysis, where quantification of Trim28 protein in RFP+ cells showed loss of Trim28 expression in the majority of neurons (83-97%) in all of the groups (Fig 2c-d).

We next queried ERV expression in adult neurons lacking Trim28. We sequenced the RNA from the isolated RFP+ nuclei and investigated the expression of ERV families as well as individual elements, using the same bioinformatical pipelines used for the NPCs. Remarkably, in contrast to the NPC experiment, we observed no activation of ERVs upon *Trim28* deletion in adult neurons (Fig 2e). We also found no transcriptional activation of any other TE classes.

These results were particularly striking since Trim28 and many of its KRAB-ZFP adaptors are expressed in the brain, suggesting it is a significant organ for Trim28-mediated TE silencing (Imbeault et al., 2017). We therefore performed a series of additional control experiments to verify this finding. To ensure that the lack of ERV activation was not due to a bystander effect of nearby glial cells in which *Trim28* could be inactivated using our experimental setup, we developed an additional CRISPR-approach for cell-type specific deletion of *Trim28* in neurons using transgenic mice that conditionally express Cas9-GFP upon Cre expression (Stop-Cas9-GFP knock-in) (Platt et al., 2014) (Fig S2a). We generated AAV-vectors expressing the gRNAs and a Cre-inducible H2B-RFP reporter and an AAV-vector expressing Cre under the control of the neuron-specific Synapsin1 (Syn) promoter. Upon transduction, Cre expression and subsequent expression of RFP and Cas9-GFP resulted in highly efficient neuron-specific gene editing of *Trim28* (Fig S2b-d). RNA-seq analysis for TE-expression revealed no ERV activation upon *Trim28* removal (Fig S2e), which were in line with our results from the ubiquitous Cas9-GFP knock-in mice. To further verify that the lack of ERV expression was not caused by potential Cas9-mediated side effects which may occur *in vivo,* we injected AAV vectors expressing Cre into the forebrain of adult floxed *Trim28* animals (Cammas et al., 2000) (Fig S2f). Again, we obtained a highly efficient *Trim28* deletion in adult mouse neurons but did not observe ERV activation (Fig S2g-i). Furthermore, ChIP-seq from adult mouse forebrain (Jiang et al., 2017) showed a lack of H3K9me3 accumulation on MMERVK10C sequences in adult neurons, in line with our observed lack of transcriptional activation upon *Trim28* deletion (Fig 2f). Taken together, these results demonstrate that Trim28 is not required to silence the transcription of ERVs in adult neurons.

### Deletion of Trim28 in NPCs *in vivo*

Our results demonstrate that Trim28 is essential for transcriptional repression of ERVs in NPCs, but not in mature neurons. This suggests the existence of an epigenetic switch, where the dynamic and reversible Trim28/H3K9me3-mediated repression found in brain development is replaced by a different stable silencing mechanism in the adult brain. This is similar to what has been observed in early development where Trim28 participates in the establishment of DNA methylation to stable silence transposable elements (Wiznerowicz et al., 2007). To test this hypothesis, we deleted *Trim28* in dividing neural progenitors *in vivo,* which give rise to mature neurons in adulthood. We bred *Emx1*-Cre transgenic mice (Iwasato et al., 2000) with *Trim28*-flox mice, resulting in a *Trim28* deletion in cortical progenitors starting from embryonic day 10 (Fig 3a, S3a). For better visualisation of *Trim28*-deleted cells by IHC we included a *Cre*-inducible GFP-reporter (gtRosa26-Stop-GFP) in the breeding scheme.

**Figure 3.**
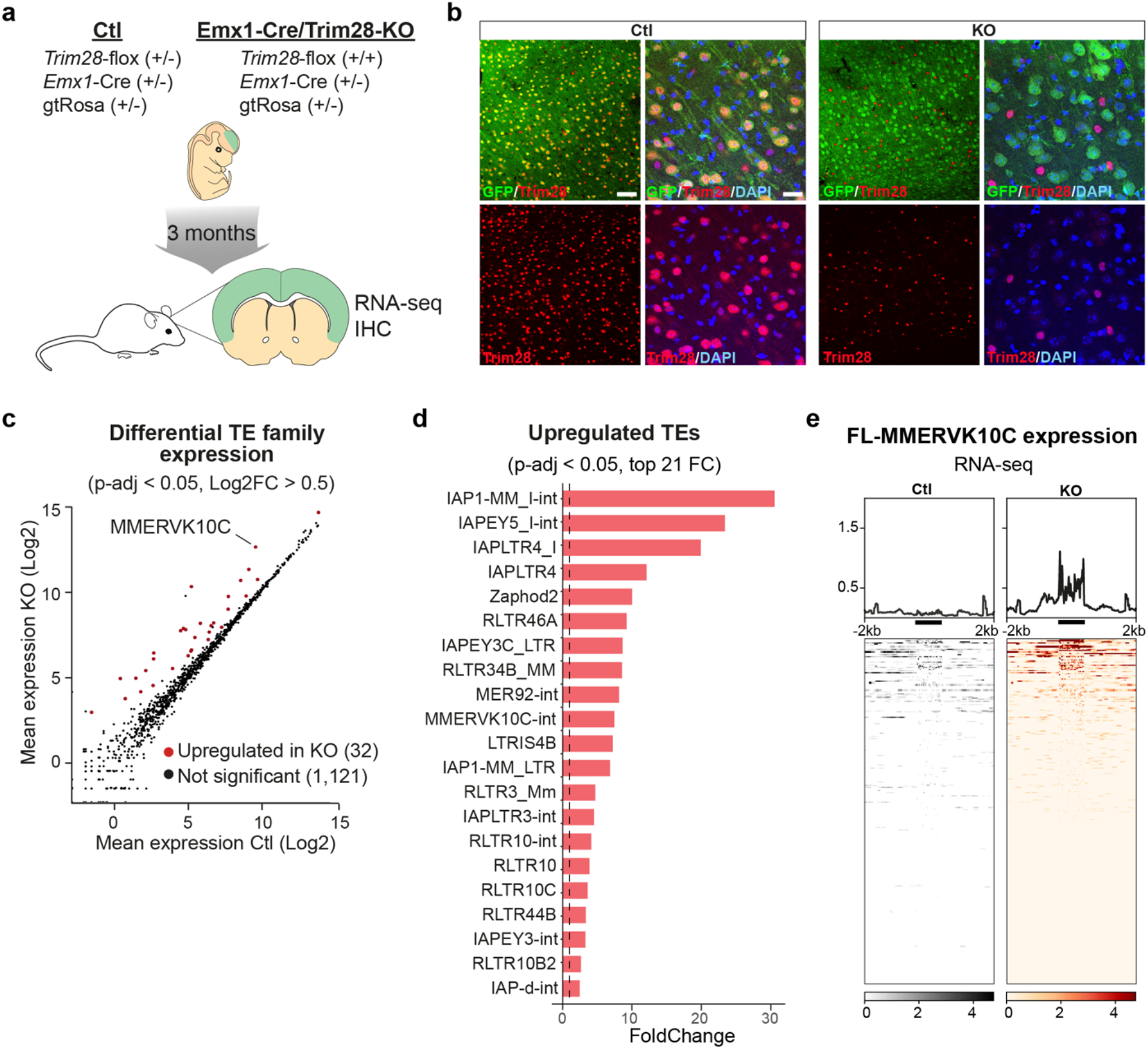
Deletion of Trim28 during brain development results in aberrant TE expression in the adult brain. (a) A schematic of the breeding scheme allowing conditional deletion of Trim28 during cortical development and analysing the adult tissue 3 months later by IHC and RNA-seq. (b) IHC for Trim28 in the adult cortex revealed that the protein was lost in the cells where Cre was expressed (GFP+ cells). Scalebars: low magnification 75 μm, high magnification 20μm. (c) RNA-seq analysis of the expression of TE-families using TEtranscripts (d) Significantly upregulated TE families upon the Trim28-KO, in which the families with the highest fold change are listed. (e) RNA-seq analysis of full length MMERVK10Cs in the adult tissue. The location of the full length MMERVK10Cs are indicated as a thick black line under each histogram.

*Emx1*-Cre (+/−), *Trim28*-flox (+/+) mice were born at the expected ratio and survived into adulthood. Their overall brain morphology and size was not affected by the loss of *Trim28* during cortical development. IHC analysis of adult brains revealed that Trim28 protein was absent in pyramidal cortical neurons and that these cells also expressed GFP, indicating an efficient Cre-mediated excision of *Trim28* during development (Fig 3b, Fig S3a). Cells that did not express GFP – for example, microglia and interneurons – expressed Trim28 as expected.

We performed RNA-seq on adult cortical tissue from these animals and control siblings to analyse ERV expression. In adult cortical tissue from *Emx1*-Cre (+/−), *Trim28*-flox (+/+) we observed a robust upregulation of several ERV families, many of which were also upregulated *in vitro* in the NPC experiment, including e.g. MMERVK10C (Fig 3c, Fig S3b – individual TEs). Similar to the NPC experiment, we found that only a few full-length elements were activated, but that these were highly expressed (Fig 3d-e). Thus, the deletion of *Trim28* in NPCs *in vivo* during brain development results in the expression of ERVs in the adult brain. Notably, and in contrast to the *Trim28*-KO in NPCs *in vitro*, we did not detect an effect on nearby gene expression by activation of ERVs (Fig S3c) nor observe any transcriptional readthrough from activated ERV elements into neighbouring genes. This also included the very same elements that had this property *in vitro* in NPCs (Fig 3e, Fig S3d). These results demonstrate that deletion of Trim28 during brain development *in vivo* results in high-level expression of ERVs in the adult brain.

### Downstream transcriptional consequences of ERV activation in the brain

Our finding that *Emx1*-Cre (+/−), *Trim28*-flox (+/+) mice survive - despite high levels of ERV expression in the brain *-* raises the question about the downstream consequences of ERV-activation *in vivo*. We first compared the expression of protein-coding genes in *Emx1*-Cre (+/−), *Trim28*-flox (+/+) mice to their control littermates. We included RNA-seq data from animals in which *Trim28* was deleted in adult neurons (AAV.Syn-Cre, *Trim28-*flox (+/+)) in this analysis since these samples provide an important control setting in which the loss of *Trim28* does not impact ERV expression but only other Trim28 targets (Fig S2f-i). Analysis of the *Emx1*-Cre (+/−), *Trim28*-flox (+/+) mice revealed 164 significantly upregulated and 86 significantly downregulated genes, of which 26 of the upregulated and 13 of the downregulated were similarly changed in the AAV-Cre, *Trim28-*flox (+/+) mice (Fig S3e-g). This indicates that the activation of ERVs during brain development causes substantial downstream effects. Gene ontology (GO) analysis on molecular and biological pathways of genes specifically altered in *Emx1*-Cre (+/−), *Trim28*-flox (+/+) - but not in AAV-Cre, *Trim28-*flox (+/+) mice - revealed significant changes in genes related to cell adhesion, indicative of synaptic functions (Fig S3h).

### Single-nuclei RNA-seq analysis of ERV-expressing brain tissue

These results indicate that the activation of ERVs in neurons results in downstream transcriptional effects that could have an impact on neuronal function. However, the brain is a complex tissue composed of several cell types located in close vicinity. To separate cell-intrinsic effects from cell-extrinsic effects, we performed single-nuclei RNA-seq analysis on forebrain cortical tissue dissected from *Emx1*-Cre (+/−), *Trim28*-flox (+/+) and control littermates. High-quality single-nuclei sequencing data was generated from a total of 14,296 cells, including 7,670 from *emx1*-Cre (+/−), *Trim28*-flox (+/+) mice and 6,626 from control littermates (Fig 4a).

**Figure 4.**
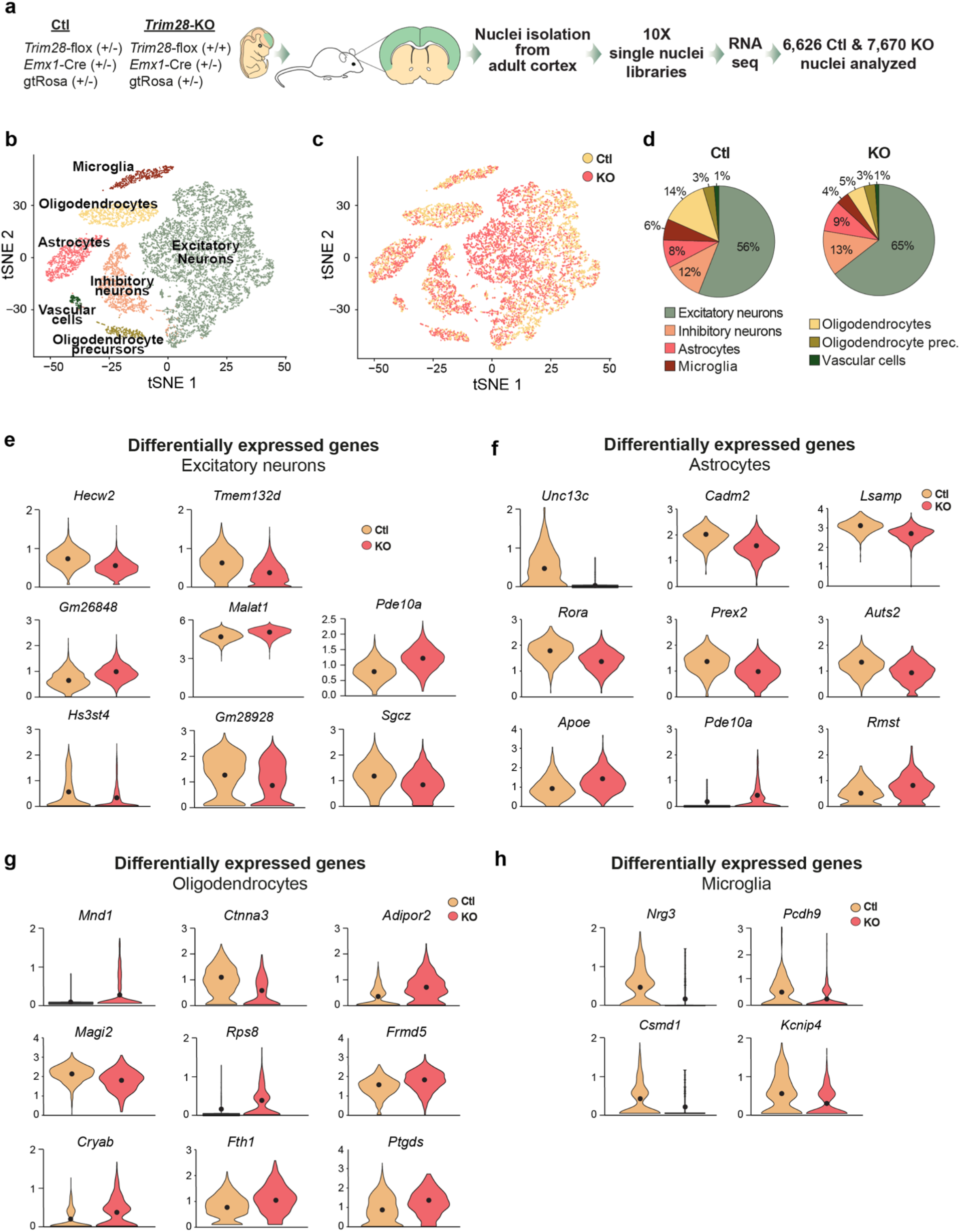
Single nuclei RNA-seq of cortical tissue from Emx1-Cre/Trim28-KO animals and their littermates. (a) A schematic of the workflow for the single nuclei RNA-seq of cortical tissue from Emx1-Cre/Trim28-KO animals and their littermates. (b) t-SNE plot showing the unbiased clustering analysis with seven different cell clusters. (c-d) t-SNE plot and pie-charts showing the distribution of Trim28-KO and control cells over the seven different clusters. There were no major differences in proportions of the different cell clusters between Emx1-Cre/Trim28-KO animals and controls. (e-h) Significant cell type specific changes in gene expression between Emx1-Cre/Trim28-KO animals and controls as revealed by single-nuclei RNA-seq. The black dots represent the mean value (p-adj value < 0.01).

We first performed an unbiased clustering analysis to identify and quantify the different cell types present in the brain tissue. We detected seven different clusters (Fig 4b, S4), including excitatory and inhibitory neurons as well as several different glial populations, with excitatory neurons making up the largest cluster with more than half of the cells (Fig 4b). Overall there was no major difference in cell number proportions between *Emx1*-Cre (+/−), *Trim28*-flox (+/+) and control animals (Fig 4c-d). For example, we found no reduction of excitatory neurons in the *Emx1*-Cre (+/−), *Trim28*-flox (+/+) animals even though this cell population lacked *Trim28.* We conclude that deletion of *Trim28* during brain development in neuronal progenitors does not result in significant cell death.

Next we analyzed transcriptional differences between the two genotypes. Among the dysregulated genes in excitatory neurons, we found altered expression of several lncRNAs and protein-coding genes. Notably, we observed reduced expression of *Hecw2*, a ubiquitin ligase linked to neurodevelopmental delay (Berko et al., 2017) and *Sgcz*, a transmembrane protein linked to mental retardation (Piovani et al., 2014) (Fig 4e). We also found transcriptional alterations in astrocytes and oligodendrocytes (Fig 4f-g), two additional cell types in which *Emx1* is expressed during brain development and therefore should lack Trim28 expression. Among the dysregulated genes in astrocytes, we detected upregulation of *Apoe*, the key risk variant gene for Alzheimer’s disease (Yamazaki et al., 2019), downregulation of *Lsamp*, a membrane located protein linked to depression and panic disorder (Koido et al., 2012), *Rora*, a nuclear hormone receptor linked to intellectual disability and epilepsy (Guissart et al., 2018) and *Auts2*, a transcriptional regulator linked to several neurodevelopmental disorders (Oksenberg and Ahituv, 2013). In oligodendrocytes we observed upregulation of *Fth1*, a ferritin gene linked to neurodegeneration (Muhoberac and Vidal, 2019). In contrast, we detected no evidence of altered gene expression in interneurons, a neuronal subtype that maintains *Trim28* expression in *Emx1*-Cre (+/−), *Trim28*-flox (+/+) mice.

Interestingly, we noted that microglial cells displayed transcriptome alterations. For example, microglia showed a reduced expression of *Nrg3*, a ligand to tyrosine kinase receptors that has been linked to schizophrenia (Kao et al., 2010) and *Csmd1*, a complement regulatory gene linked to familial epilepsy (Naseer et al., 2016). Microglia are immune cells of endodermal origin that do not express Emx1 during development and should therefore maintain *Trim28* expression in *Emx1*-Cre (+/−), *Trim28*-flox (+/+) animals. Taken together, these results therefore suggest that the ERV activation in excitatory neurons, due to the loss of Trim28 during brain development, results in both cell-autonomous and non-cell autonomous effects, specifically on microglia where some of the downstream dysregulated genes have been previously linked to psychiatric disorders.

### Cell type specific analysis of ERV activation in the brain

The single-nuclei RNA-seq analysis indicated that microglia display transcriptional alterations despite maintaining Trim28 expression. Thus, microglia should be affected through cell-extrinsic mechanism mediated by derepressed ERVs from adjacent cells lacking Trim28. To verify this observation, we devised a strategy to analyze the expression of ERVs in the different cell populations in the *Emx1*-Cre (+/−), *Trim28*-flox (+/+) animals and control littermates. Since current pipelines available for high-throughput single-cell RNA-seq analysis do not allow for estimation of TE-expression, we developed TrustTEr: a new bioinformatical pipeline that initially uses the cell clusters established based on the expression of protein-coding genes (Fig 4b). By back-tracing the reads from cells forming each cluster we were then able to analyze the expression of ERVs and other TEs using TEtranscripts in distinct cell populations (Fig 5a) [21], increasing the sensitivity of TE expression at single cell type resolution.

**Figure 5.**
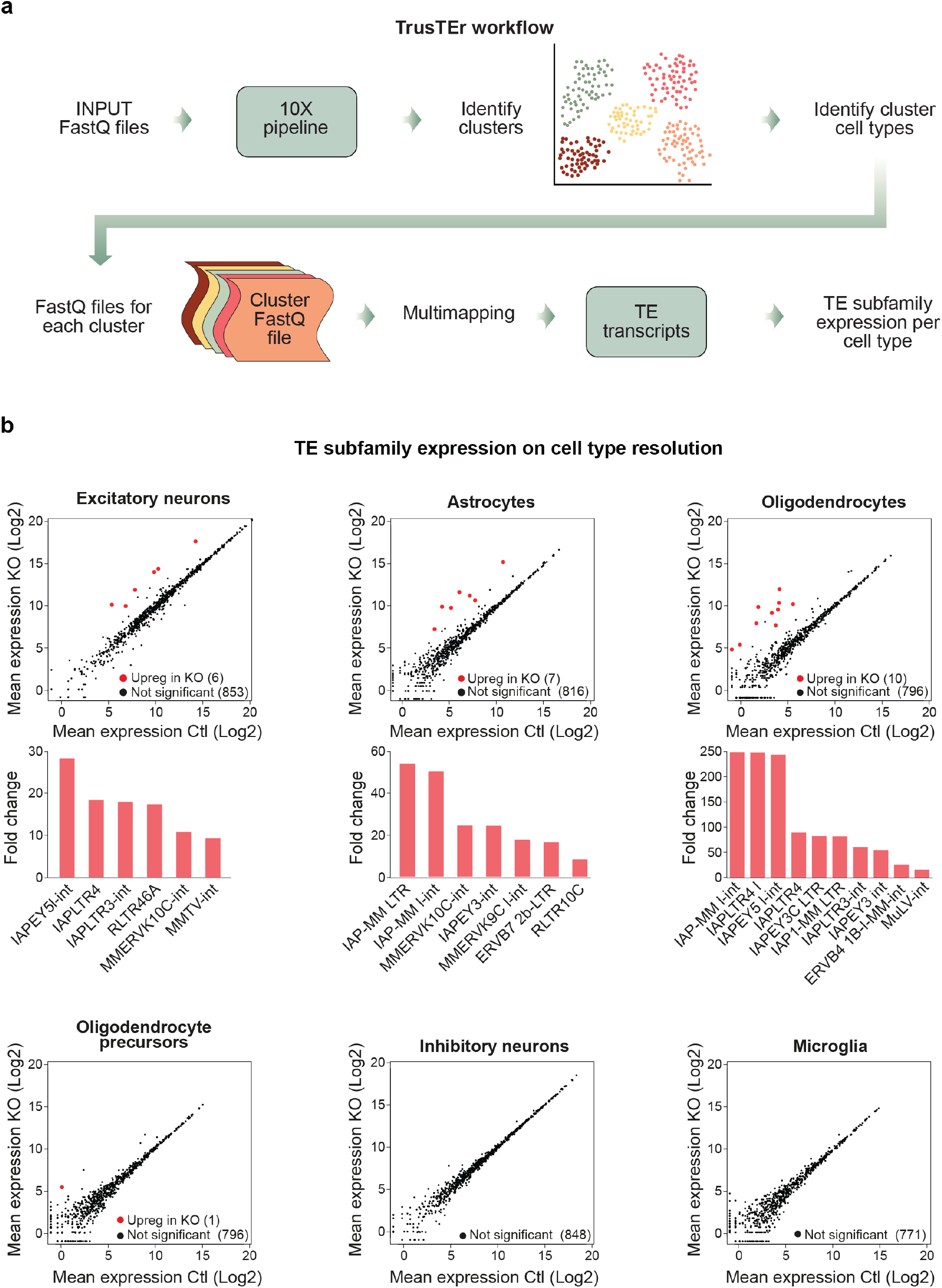
Cell type specific analysis of ERV activation in Emx1-Cre/Trim28-KO animals. (a) TrusTEr: an approach to analyse ERV expression in single nuclei RNA-seq data. (b) Mean plots show the changes of TE subfamily expression in each cell cluster upon the Emx1-Cre/Trim28-KO. TEs were upregulated in cell types in which Trim28 was deleted (excitatory neurons, astrocytes, oligodendrocytes and oligodendrocyte precursors). The specific upregulated elements and their fold changes are listed in bar graphs under each mean plot. In cell types in which Trim28 was not deleted, TE expression remained unaffected (inhibitory neurons and microglia).

Using TrusTEr we found that different ERV families (e.g. MMERVK10C) were expressed in excitatory neurons, astrocytes, oligodendrocytes and oligodendrocytes progenitors i.e. the cell types in which *Emx1* is expressed during brain development and thus lack Trim28 (Fig 5b). Notably, we found differences in ERV expression in distinct cell-types, including activation of different families, demonstrating a cell-type specificity in the response to *Trim28* deletion. However, we found no upregulation of ERV expression in interneurons and microglia where Trim28 is maintained (Fig 5b) This analysis confirms that the transcriptional alterations observed in microglia are due to downstream effects of ERV activation in surrounding cell types.

### Expression of ERVs in the brain is linked to inflammation

The microglia response to *Trim28*-KO in neurons as intriguing as the aberrant expression of ERVs and other TEs have been linked to inflammation (Hurst and Magiorkinis, 2015; Ishak et al., 2018; Lim et al., 2015; Roulois et al., 2015; Saleh et al., 2019; Tam et al., 2019a; Thomas et al., 2017). To study this further, we performed IHC analysis followed by specific analysis of microglia after labelling of inflammatory factor Iba1. We found that the microglial cells in *Emx1*-Cre (+/−), *Trim28*-flox (+/+) mice displayed clear signs of activation (Fig 6a). These microglia expressed higher levels of Iba1 and displayed a morphology that is typical for a mild activation phenotype, including longer and thicker processes with increased numbers of branches (Fig 6b). Interestingly, we only found activated microglia in the cortex, where excitatory neurons lack Trim28, but not in the nearby forebrain structure striatum, where Trim28 is still expressed in all neurons (Fig 6c). The neuroinflammation was therefore spatially restricted to the area with increased ERV expression. In addition, we found no signs of neuroinflammation in animals where *Trim28* was deleted in mature neurons, a setting where Trim28 is removed but no ERVs are activated (AAV.Syn-Cre/*Trim28-*flox) (Fig S5a). These results verify that neuroinflammation was not activated by the loss of *Trim28 per se* or by direct *Trim28*-targets in neurons, but rather by the expression of ERVs.

**Figure 6.**
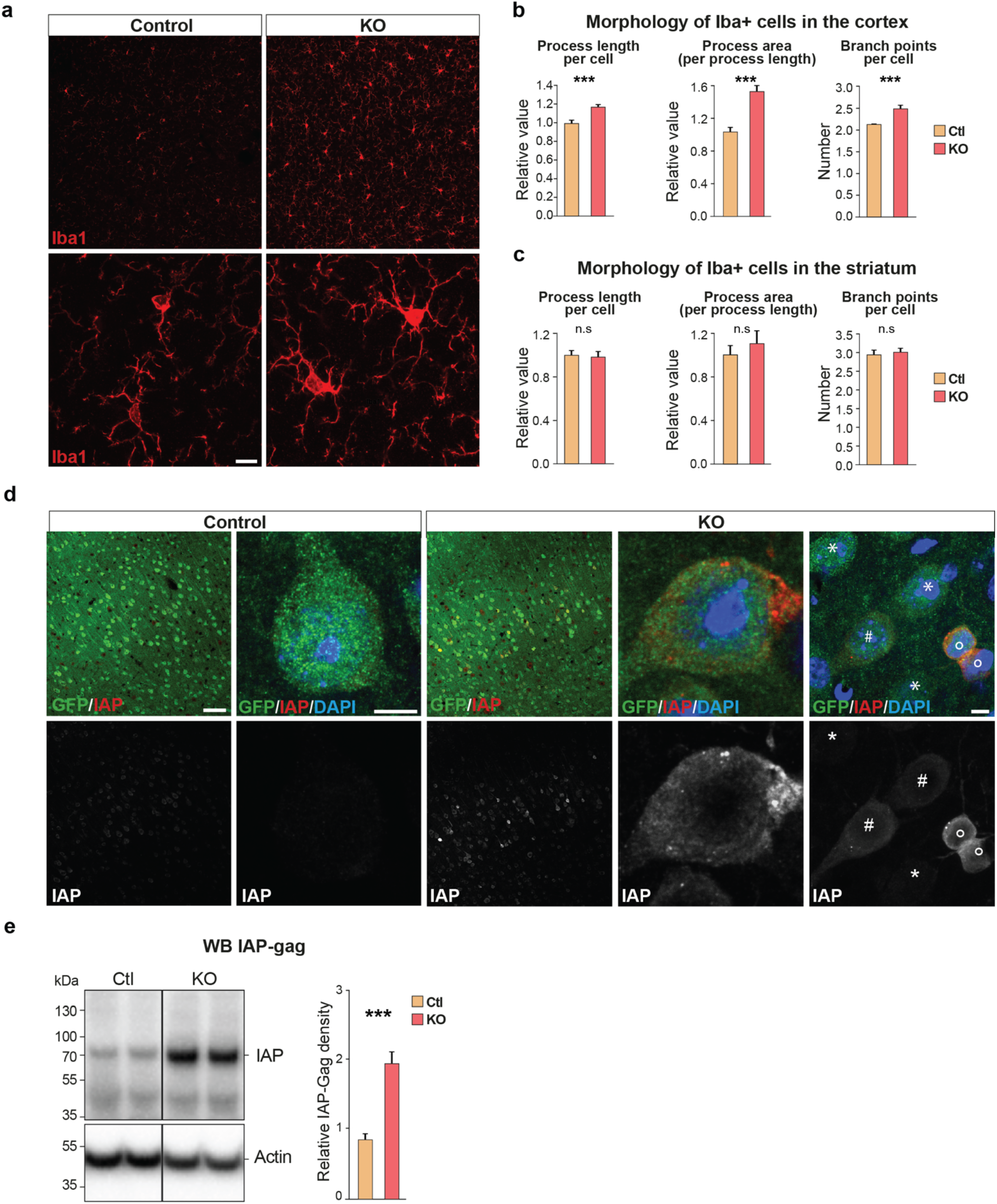
Activation of ERVs during brain development results in the presence of ERV proteins in adult brain tissue and signs of neuroinflammation. (a) IHC analysis for the microglia marker in Emx1-Cre/Trim28-KO animals and controls. Scalebars: Low magnification 75 μm, high magnification 10 μm (b-c) The morphology of microglia in cortex and striatum (control) of Emx1-Cre/Trim28-KO animals and controls were quantified by high content screening of Iba1 immunoreactivity, revealing differences in process length, area and number of branch points specifically in cortex (d) Immunohistochemistry for IAP-gag visualized the presence of ERV proteins in the cortex of Emx1-Cre/Trim28-KO animals. The distribution of IAP-gag was heterogenous among Trim28-KO neurons, where it was either accumulated in a low (#) or high (°) number of aggregates in the cytoplasm, or as weak homogenous staining throughout the cytoplasm (#) or not present at all (*) Scalebars: low magnification 75μm, high magnification 5 μm. (e) Western blot for IAP-gag of cortical tissue from adult Emx1-Cre/Trim28-KO animals (n=4) and control littermates (n=5). ***p-value < 0.001

### Aggregates of ERV-derived proteins are found in areas of neuroinflammation

A number of recent studies have demonstrated that the presence of ERV-derived nucleic acids can activate the innate immune system, such as double-stranded RNAs or single stranded DNA (Hurst and Magiorkinis, 2015; Ishak et al., 2018; Roulois et al., 2015). According to this model the host cells misinterpret the expression of ERVs as a viral infection, and this triggers an autoimmune response through the activation of an interferon signaling. Therefore, we searched for evidence of an interferon response and activation of a viral defense pathway in the transcriptome data (bulk RNA-seq and single nuclei RNA-seq) from *Emx1*-Cre (+/−), *Trim28*-flox (+/+) mice. However, we found no evidence of activation of these pathways suggesting that the expression of genes linked to the innate immune response and viral response were not transcriptionally activated despite high levels of ERVs in the brain (Fig S5 b-c). Similarly, we found no evidence of this response in the *Trim28*-KO NPCs *in vitro*, in which ERVs were activated. Thus, deletion of *Trim28* and subsequent ERV activation in neural cells does not result in a detectable interferon response, suggesting that other mechanisms are responsible for triggering the observed neuroinflammation.

An alternative mechanism for ERV-mediated triggering of neuroinflammation is the expression of ERV-derived peptides and proteins, which could have neurotoxic properties (Li et al., 2015). To search for evidence of ERV-derived proteins, we performed WB and IHC analysis using an antibody detecting IAP-Gag protein in the brain of *Emx1*-Cre (+/−), *Trim28*-flox (+/+) mice. We found high levels of IAP-Gag protein expression in the brain of mice expressing elevated ERV transcripts, demonstrating their efficient translation into proteins (Fig 6d-e). The IAP-Gag labelling was restricted to cortical excitatory neurons lacking Trim28, as visualized by the co-expression of the Cre-dependent GFP reporter. Notably, the IAP-Gag labelling was not uniform, as some neurons expressed higher levels of IAP-Gag and we found clear evidence of ERV-derived protein aggregation (Fig 6d), suggesting that the expression of ERVs in the brain is associated with aggregation of ERV-derived proteins, which could be the underlying reason of the apparent neuroinflammation.

## Discussion

In this study we define epigenetic mechanisms that control the expression of ERVs during brain development and investigate the consequences of their inactivation. Previous work has implicated ERVs and other TEs in several neurological disorders, such as MS, AD, ALS, PD and schizophrenia, where an increased expression of TEs has been reported along with the speculation of their contribution to the disease process (Andrews et al., 2000; Douville et al., 2011; Garson et al., 1998; Guo et al., 2018; Jonsson et al., 2020; Karlsson et al., 2001; Li et al., 2015; MacGowan et al., 2007; Perron et al., 1997; Perron et al., 2008; Steele et al., 2005; Sun et al., 2018; Tam et al., 2019b). However, these studies have been difficult to interpret since the clinical observations were correlative and since the modeling of ERV activation in an experimental setting is challenging. Thus, the putative involvement of ERVs in brain disorders has remained controversial and direct experimental evidence on the consequences of ERV activation in the brain has been needed (Jonsson et al., 2020; Tam et al., 2019a). This study provides direct *in vivo* evidence that aberrant activation of ERVs during brain development triggers neuroinflammation linked to ERV-derived protein aggregation in the adult brain. This increases our understanding of the consequences of ERV activation in the brain and provides new mechanistic insights opening up for further research into the role of ERVs and other TEs in various brain disorders.

Our results demonstrate that brain development is a critical period for the repression of ERVs. During this period, Trim28 dynamically controls the expression of ERVs through a mechanism linked to the repressive histone mark H3K9me3. Remarkably, Trim28 is also required for the establishment of a different, stable silencing mechanism that is present in adult neurons – most likely DNA methylation (Wiznerowicz et al., 2007). Importantly, once established this system is independent of Trim28 and H3K9me3. This observation suggests that the ERV silencing machinery is particularly vulnerable during brain development and that perturbation of the system during this period could have life-long consequences. If Trim28-mediated silencing of ERVs is impaired during brain development, for example through mutations in KRAB-ZFPs or by environmental influence, ERVs may be derepressed in adult neurons, resulting in neuroinflammation.

It is well established that many psychiatric disorders are a consequence of neurodevelopmental alterations (Bale et al., 2010; Horwitz et al., 2019). A combination of genetic components and environmental exposures are thought to contribute to the appearance and progress of these diseases, indicating that epigenetic alterations play a key role in the disease process (see e.g. (Khashan et al., 2008; Susser et al., 2008)). Many psychiatric disorders also have an immune component that is thought to play an important role in the disease process (Bauer and Teixeira, 2019). This immune response includes both peripheral activation as well as glial activation, in the brain. The underlying cause for this immune response remains largely unknown. However, the combination of epigenetic dysregulation and immune activation in psychiatric disorders, together with the observations made in the current study, suggest that ERVs are directly involved in the disease process. We have previously demonstrated that the deletion of *Trim28* during postnatal forebrain development or heterozygous deletion of *Trim28* during early brain development results in complex behavioral changes including hyperactivity and an impaired stress-response (Jakobsson et al., 2008) (Fasching et al., 2015). In addition, heterozygous germ line deletion of *Trim28* in mice has been described to provoke an abnormal exploratory behavior (Whitelaw et al., 2010). Together, these findings demonstrate that disruption of Trim28 levels in the mouse brain results in behavioral changes, similar to impairments found in psychiatric disorders. The current study links the deletion of *Trim28* and the disease-related phenotypes to ERV activation.

In the current study we found no evidence that activation of ERVs in NPCs, *in vitro* or *in vivo*, triggers an interferon response. Rather, our results point to an important role for ERV-derived peptides and proteins in the observed neuroinflammation. We found numerous neurons displaying expression of ERV-derived proteins in the transgenic mice that expressed high levels of ERVs. These proteins were not distributed in a uniform pattern, but rather tended to aggregate in a subset of neurons often located in close vicinity. This observation is interesting given the well-established link between protein aggregation and neurodegenerative disorders (Ross and Poirier, 2004). It is plausible that the ERV-protein aggregates leads to neuroinflammation and may also serve as a trigger for further protein aggregation, although further in-depth studies are needed to understand this phenomenon.

In summary, these results demonstrate that Trim28 is required to silence ERVs during brain development and that the perturbation of this system results in ERV-mediated neuroinflammation in the adult brain. These results provide a new perspective to the potential cause and progression of neurodevelopmental and neurodegenerative disorders and further research into ERV-dysregulation in these types of disorders is therefore warranted.

## Methods

### Generation of Cas9-GFP mouse NPC cultures

The forebrain was dissected on embryonic day 13.5 from embryos obtained by breeding homozygote Cas9-GFP knock-in mice (Platt et al., 2014). The tissue was mechanically dissociated and plated in gelatin coated flasks and maintained as a monolayer culture (Conti et al., 2005) in NSA medium (Euromed, Euroclone) supplemented with N2 hormone mix, EGF (20 ng/ml; Gibco), bFGF (20 ng/ml; Gibco), 2nM L-glutamine and 100ug/ml Pen/Strep. Cells were then passaged 1:3-1:6 every 2-3 days using Accutase (Gibco).

### Targeting *Trim28 in vitro*

Guides were designed at crispr.mit.edu and are listed in supplementary table 1. Lentiviruses were produced according to (Zufferey et al., 1997) and titers were 10^9^ TU/ml, which was determined using qRT-PCR. Cas9 mouse NPCs where transduced at a MOI 40 and allowed to expand for 10 days prior to FACS (FACSAria, BD Biosciences). Cells were detached and resuspended in basic culture media (media excluding growth factors) with propidium iodide (BD Biosciences) and strained (70μm filters, BD Biosciences). RFP cells were FACS isolated at 4°C (reanalysis showed >99% purity) and pelleted at 1500 rpm for 5 min, snap frozen on dry ice and stored at −80°C until RNA/DNA were isolated. All groups were performed in biological triplicates.

### Targeting *Trim28 in vivo* using CRISPR/Cas9 in the adult brain

All animal-related procedures were approved by and conducted in accordance with the committee for use of laboratory animals at Lund University.

The production of AAV5 vectors has been described in detail elsewhere (Ulusoy et al., 2009) and titers were in the order of 10^13^ TU/ml, which was determined by qRT-PCR using taqman primers towards the ITR. Prior to injection, the vectors were diluted in PBS; the vectors containing the guide RNAs were diluted to 30% except upon co-injection of guides 3, 4, and 13 where the vectors were diluted to 10% each. Rosa26 Cas9 knock-in mice where anesthetized by isoflurane prior to the intra-striatal injections (coordinates from bregma: AP +0.9 mm, ML +1.8 mm, DV −2.7 mm) of 1 μl virus solution (0.1 μl / 15 s). The needle was kept in place for additional 2 minutes post injection to avoid backflow. Animals were sacrificed after two months and analyzed either by IHC or nuclei isolation (see details below) followed by DNA- or RNA-seq.

### Targeting *Trim28 in vivo* during neural development

Male *Emx1*-Cre (+/−); *Trim28*-fl (+/−); gtRosa(+/−) were bred with *Trim28*-fl (+/+) females to generate animals in which one (*Emx1*-Cre +/−; *Trim28*-fl+/−) or both (*Emx1*-Cre +/−; *Trim28*-fl +/+) *Trim28* alleles had been excised, used as control and *Trim28*-KO, respectively. Animals used for IHC were additionally heterozygote for gtRosa, in order to visualize the cells in which Cre had been expressed. Animals were genotyped from tail biopsies according to previous protocols (Cammas et al., 2000) and sacrificed at 3 months of age for either IHC or RNA sequencing.

### Immunohistochemistry

Mice were given a lethal dose of pentobarbiturate and transcardially perfused with 4% paraformaldehyde (PFA, Sigma), the brains were post-fixed for two hours and transferred to phosphate buffered saline (PBS) with 25% sucrose. Brains were coronally sectioned on a microtome (30μm) and put in KPBS. IHC was performed as described in detail elsewhere (Sachdeva et al., 2010). Antibodies: Trim28 (Millipore, 1:500), IAP-gag (a kind gift from Bryan Cullen and described in (Dewannieux et al., 2004), 1:2000), Gfap (DAKO; 1:1000), Iba1 (WAKO, 1:1000). All sections were counterstained with 4’,6-diamidino-2-phenylindole (DAPI, Sigma-Aldrich, 1:1000). Secondary antibodies from Jackson Laboratories were used at 1:400.

### Nuclei isolation

Animals were sacrificed by cervical dislocation and brains quickly removed. The desired regions were dissected and snap frozen on dry ice and stored at −80°C. The nuclei isolation was performed according to (Sodersten et al., 2018). In brief, the tissue was thawed and dissociated in ice-cold lysis buffer (0.32 M sucrose, 5 mM CaCl_2,_ 3 mM MgAc, 0.1 mM Na_2_EDTA, 10 mM Tris-HCl, pH 8.0, 1 mM DTT) using a 1 ml tissue douncer (Wheaton). The homogenate was carefully layered on top of a sucrose cushion (1.8 M sucrose, 3 mM MgAc, 10 mM Tris-HCl, pH 8.0, and 1 mM DTT) before centrifugation at 30,000 × *g* for 2 hours and 15 min. Pelleted nuclei were softened for 10 min in 100 μl of nuclear storage buffer (15% sucrose, 10 mM Tris-HCl, pH 7.2, 70 mM KCl, and 2 mM MgCl_2_) before resuspended in 300 μl of dilution buffer (10 mM Tris-HCl, pH 7.2, 70 mM KCl, and 2 mM MgCl_2_) and run through a cell strainer (70 μm). Cells were run through the FACS (FACS Aria, BD Biosciences) at 4°C with low flowrate using a 100 μm nozzle (reanalysis showed >99% purity). Sorted nuclei intended for either DNA or RNA sequencing were pelleted at 1,300 x *g* for 15 min and snap frozen, while nuclei intended for single nuclei RNA sequencing were directly loaded onto the 10X Genomics Single Cell 3’ Chip – see *Single nuclei sequencing*.

### Analysis of CRISPR/Cas9 mediated *Trim28*-indels

Total genomic DNA was extracted from all *Trim28*-KO and control groups using DNeasy blood and tissue kit (Qiagen) and a 1.5 kb fragment surrounding the different target sequences were amplified by PCR (primers can be found in supplementary table 2) before subjected to NexteraXT fragmentation, according to manufacturer recommendations. Indexed tagmentation libraries were sequenced with 2×150 bp PE reads and analyzed using an in-house TIGERq pipeline to evaluate CRISPR-Cas9 editing efficiency.

### RNA-sequencing

Total RNA was isolated from frozen cell/nuclei pellets and brain tissue using the RNeasy Mini Kit (Qiagen) and used for RNA-seq and qPCR (tissue pieces were run in the tissue lyser for 2 min, 30 Hz, prior to RNA isolation). Libraries were generated using Illumina TruSeq Stranded mRNA library prep kit (poly-A selection) and sequenced on a NextSeq500 (PE 2×150bp).

The reads were mapped with STAR (2.6.0c) (Dobin et al., 2013), using gencode mouse annotation GRCm38.p6 vM19 as a guide. Reads were allowed to map to 100 loci with 200 anchors, as recommended by (Jin et al., 2015) to run TEtranscripts.

Read quantification was performed with TEtranscripts version 2.0.3 in multimode using gencode annotation GRCm38.p6 vM19 for gene annotation, as well as the curated GTF file of TEs given by TEtranscripts authors (Jin et al., 2015). The output matrix was then divided between TE subfamilies and genes to perform differential expression analysis (DEA) with DESeq2 (version 1.22.2) (Love et al., 2014) contrasting *Trim28*-KO against control samples.

Samples from the different guides targeting *Trim28* were pooled together and tested against the LacZ controls. The data was normalized using sizeFactors from the DESeq2 object (median ratio method described in (Anders and Huber, 2010) to account for any differences in sequencing depth.

In order to define differentially expressed elements and study their effects on gene expression, reads were uniquely mapped with STAR (2.6.0c). Full length ERVK predictions were done with REannotate (Pereira, 2008) and read quantification of them was performed using featureCounts (Subread 1.6.3) (Liao et al., 2014). Differential expression analysis (DEA) was done with DESeq2. An intersection of the gencode annotation GRCm38.p6 vM19 with windows of 10, 20 and 50 kbp up and downstream of the upregulated elements was made with BEDtools intersect (Quinlan and Hall, 2010), same intersection was done for non-upregulated elements to compare their nearby gene dysregulation.

Bigwig files were normalized by RPKM using bamCoverage from deeptools and uploaded to USCS Genome Browser (release GCF_000001635.25_GRCm38.p5 (2017-08-04)).

### Single nuclei RNA-sequencing

Nuclei were isolated from the cortex of *Emx1*-Cre(+/−)/*Trim28*-fl(+/−,+/+) animals (Ctl n=2, KO n=2) as described above. 8500 nuclei per sample were sorted via FACS and loaded onto 10X Genomics Single Cell 3’ Chip along with the reverse transcription mastermix following the manufacturer’s protocol for the Chromium Single Cell 3′ Library (10X Genomics, PN-120233) to generate single-cell gel beads in emulsion. cDNA amplification was done as per the guidelines from 10x Genomics and sequencing libraries were generated with unique sample indices (SI) for each sample. Libraries for samples were multiplexed and sequenced on a Novaseq using a 150-cycle kit.

The raw base calls were demultiplexed and converted to sample specific fastq files using cellranger mkfastq^1^ that uses bcl2fastq program provided by Illumina. The default setting for bcl2fastq program was used, allowing 1 mismatch in the index, and raw quality of reads was checked using FastQC and multiQC tools. For each sample, fastq files were processed independently using cellranger count version 3.0 pipeline (default settings). This pipeline uses splice-aware program STAR^5^ to map cDNA reads to the transcriptome (mm10). Since in nuclei samples it is expected a to get a higher fraction of pre-mRNA, a pre-mRNA reference was generated using cellranger guidelines.

Mapped reads were characterized into exonic, intronic and intergenic if at least 50% of the read intersects with an exon, intronic if it is non-exonic and it intersects with an intron and intergenic otherwise. Only exonic reads that uniquely mapped to transcriptome (and the same strand) were used for the downstream analysis.

Low quality cells were filtered out based on fraction of total number of genes detected in each cell (±3 nmads). From the remaining 14,296 nuclei,6,626 came from control samples (Ctl) and 7,670 from knock-out (KO).

For downstream analysis, samples were merged together using Seurat (version 3) R package (Dobin et al., 2013). Clusters have been defined with Seurat function FindClusters using resolution 0.1 and visualized with t-SNE plots. Cell type annotation was performed using both known marker-based expression per cluster and a comparison of the expression profiles of a mouse brain Atlas (Zeisel et al., 2018). A marker gene set consisting of up-regulated gene per cluster among the cells, combined with marker genes for all the 256 cell types in the atlas was used in the comparison. The 256 atlas cell types were grouped into main clusters at Taxonomy rank 4 (39 groups) and mean expression per group was calculated using the marker gene set. These were compared to the mean expression in our clusters using Spearman correlation. Based on clusters annotation, clusters 0, 1, 2, 5 and 6, 7 were manually merged as excitatory and inhibitory neurons respectively. Cell barcodes for all clusters were extracted and original .bam files obtained from the cellranger pipeline were used to subset aligned files for each cluster (subset-bam tool provided by 10X). Each .bam file was then converted back to clusters’ fastq files using bamtofastq tool from 10X.

The resulting fastq files were mapped using default parameters in STAR using gencode mouse annotation GRCm38.p6 vM19 as a guide. The resulting bam files were used to quantify reads mapping to genes with featureCounts (forward strandness). The output matrix was then used to calculate sizeFactors with DESeq2 that would later be used to normalize TE counts.

The cluster fastq files were also mapped allowing for 100 loci and 200 anchors, as recommended by TEtranscripts authors. Read quantification was then performed with TEtranscripts in multi-mode (forward strand) using GRCm38.p6 vM19 for the gene annotation, and a curated GTF file of TEs given by TEtranscripts’ authors.

For the data presented in figure 5b, the fold change bar plots were made from a DEA performed with DESeq2 of TE subfamilies of each cell type comparing control and knock out samples. The mean plots in the same figure, were normalized using the sizeFactors resulting from the gene quantification with the default parameters’ mapping.

### CUT&RUN

The CUT&RUN were performed according to (Skene and Henikoff, 2017). In brief, 200.000 mouse NPCs were washed, permeabilized and attached to ConA-coated magnetic beads (Bang Laboratories) before incubated with the H3K9me3 (1:50, ab8898, Abcam) antibody at 4°C overnight. Cells were washed and incubated with pA-MNase fusion protein and digestion was activated by adding CaCl_2_ at 0°C. The digestion was stopped after 30 min and the target chromatin released from the insoluble nuclear chromatin before extracting the DNA. Experimental success was evaluated by capillary electrophoresis (Agilent) and the presence of nucleosome ladders for H3K9me3 but not for IgG controls.

The library preparation was performed using the Hyper prep kit (KAPA biosystems) and sequenced on NextSeq500 2×75 bp. Mapping of the reads to mm10 was performed with Bowtie2 2.3.4.2 (Langmead and Salzberg, 2012) using default settings for local alignment. Multi mapper reads were filtered by SAMtools view version 1.4 (Li et al., 2009).

Using the ERVK prediction described in the section *RNA-sequencing*, we retrieved full length MMERVK10Cs. An ERVK was considered to be a full length MMERVK10C when an annotated MMERVK10C-int of mm10 RepeatMasker annotation (open-4.0.5 – Repeat Library 20140131) would overlap more than 50% into the full length ERVK prediction. The intersection was performed with BEDtools intersect 2.26.0 (-f 0.5) (Quinlan and Hall, 2010). Heatmaps and profile plots were produced using deepTools’ plotHeatmap (Ramirez et al., 2016) and sorted using maximum expression of the *Trim28*-KO samples or guide 3 for the *in vitro* and *in vivo* CRISPRs. Tracks for genome browser were normalized using RPKM using deepTools’ bamCoverage (version 2.4.3) (Ramirez et al., 2016).

The H3K9me3 ChIPseq data from adult mouse cortex was retrieved from (Jiang et al., 2017), mapped, and analyzed in the same way as the in-house Cut & Run samples described above.

### Western blot

Dissected cortical pieces from the *Emx1*-Cre(+/−); *Trim28*-fl(+/− and +/+) animals (Ctl n=5, KO n=5) were put in RIPA buffer (Sigma-Aldrich) containing Protease inhibitor cocktail (PIC, Complete, 1:25) and lysed at 4°C using a TissueLyser LT (Qiagen) on 50 Hz for 2 min, twice, and then kept on ice for 30 min before spun at 10,000 x g for 10 min at 4°C. Supernatants were collected and transferred to a new tube and stored at −20°C. Each sample was mixed 1:1 (10 μl + 10 μl) with Laemmli buffer (BioRad) and boiled at 95°C for 5 min before loaded onto a 4-12% SDS/PAGE gel and run at 200 V before electrotransferred to a membrane using Transblot-Turbo Transfer system (BioRad). The membrane was then washed 2 × 15 min in TBS with 0.1% Tween20 (TBST), blocked for 1 hour in TBST containing 5% non-fat dry milk and then incubated at 4°C overnight with rabbit anti-IAPgag (1:10,000, a kind gift from Bryan Cullen and described in (Dewannieux et al., 2004)) diluted in TBST with 5% non-fat dry milk. The membrane was washed in TBST 2 × 15 min and incubated for 1 hour in room temperature with HRP-conjugated anti-rabbit antibody (Sigma-Aldrich, NA9043, 1:2,500) diluted in TBST with 5 % non-fat dry milk. The membrane was washed 2 × 15 min in TBST again and 1 × 15 min in TBS, before the protein expression was revealed by chemiluminescence using Immobilon Western (Millipore) and the signal detected using a Chemidoc MP system (BioRad). The membrane was stripped by treating it with methanol for 15 s followed 15 min in TBST before incubating it in stripping buffer (100 mM 2-mercaptoethanol, 2% (w/v) SDS, 62.4 mM Tris-HCL, pH 6.8) for 30 min 50°C. The membrane was washed in running water for 15 min, followed by 3 × 15 min in TBST before blocked for 1 hour in TBST containing 5% non-fat dry milk. The membrane was then stained and visualised for β-actin (rabbit anti-β-actin HRP, Sigma-Aldrich, A3854, 1:50,000) as described above.

### Morphological analysis

The morphology of Iba1+ cells in the Emx1-Cre/Trim28-fl animals (Ctl n = 3, KO n=2) was analysed through an unbiased, automated process using the Cellomics Array Scan (Array Scan VTI, Thermo Fischer). The scanner took a high number of photos (20x) throughout cortex (Ctl n = 361, KO n = 215) and striatum (Ctl n = 104, KO n = 58) and the program “Neuronal profiling” allowed analysis of process length, process area and branchpoints per cell.

## Code availability

The pipeline, configuration files and downstream analyses are available in the src folder at GitHub (https://github.com/ra7555ga-s/trim28_Jonsson2020.git). All downstream analysis and visualization were performed in R 3.5.1.

## Data availability

There are no restrictions in data availability. Accession code for the RNA and DNA sequencing data presented in this study is XXXXXX.

## Acknowledgements

We would like to thank Molly Gale Hammell, Magdalena Götz, Sten Linnarsson, Chris Douse, Florence Cammas and Bryan Cullen for providing valuable reagents and input on the manuscript. We also thank, M. Persson Vejgården, U. Jarl, and A. Hammarberg for technical assistance. We are grateful to all members of the Jakobsson lab. The work was supported by grants from the Swedish Research Council (2018-02694, JJ & 2018-03017, PJ), the Swedish Brain Foundation (FO2019-0098, JJ), Cancerfonden (190326, JJ), Barncancerfonden (PR2017-0053, JJ), Formas (2018-01008, PJ) and the Swedish Government Initiative for Strategic Research Areas (MultiPark & StemTherapy).

## Author Contributions

All authors took part in designing the study as well as interpreting the data. M.E.J. and J.J. conceived and designed the study. M.E.J., R.P., J.G.J., P.A.J, D.A.M.A., K.P., S.M., D.Y., R.R. performed experimental research and R.G, P.J. and Y.S. performed bioinformatical analyses, J.L. E.S., J.H. contributed resources. M.E.J., R.G., and J.J. wrote the manuscript and all authors reviewed the final version.

## Supplemental Information

**Supplementary figure 1.**
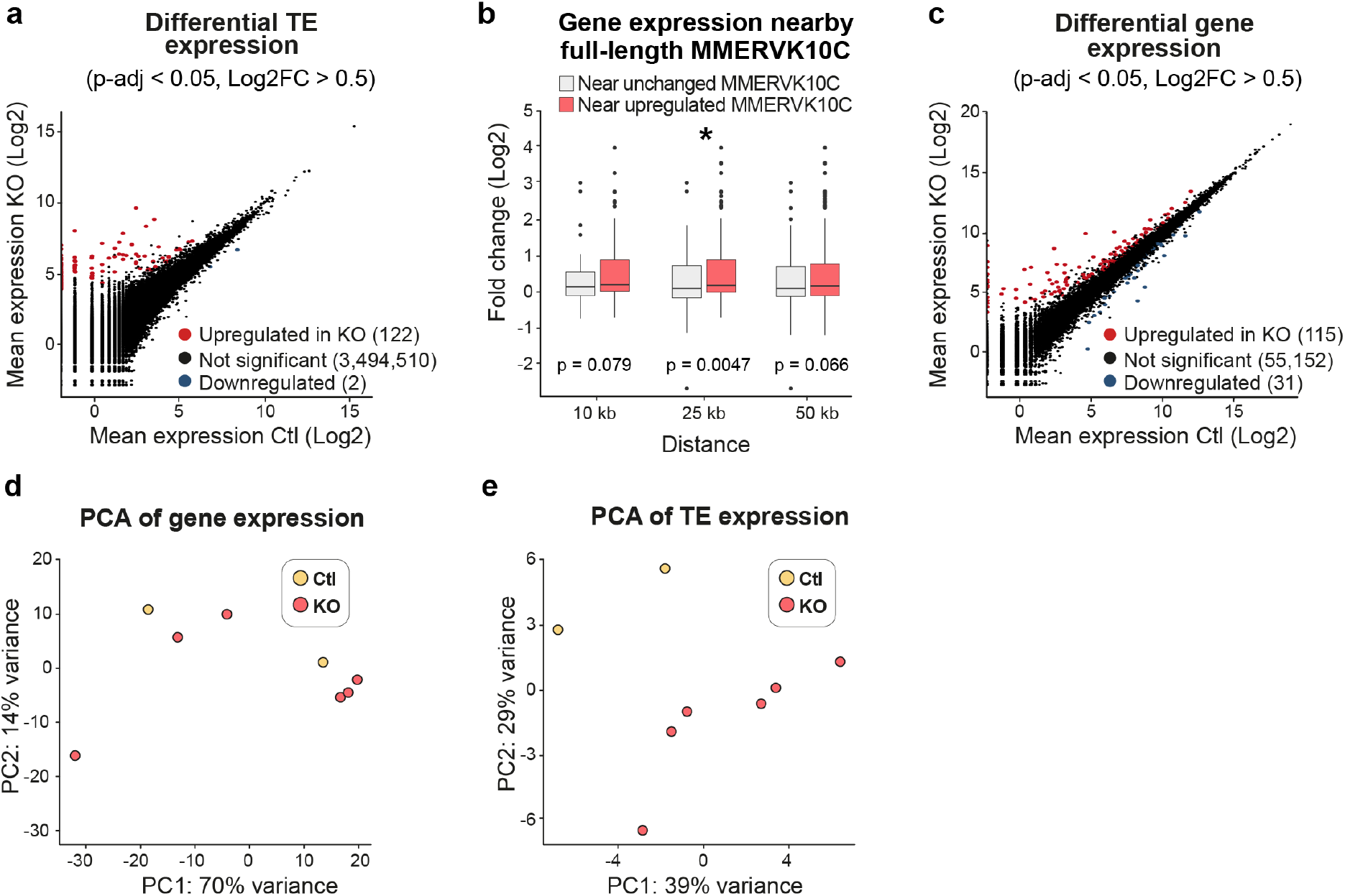
(a) RNA-seq of individual TEs in mouse NPCs upon CRISPR/Cas9-mediatied disruption of Trim28. (b) Genes located in the close vicinity of upregulated MMERVK10C elements were significantly upregulated. x-axis indicate the window-of-inclusion for genes located close to an MMERVK10C. (c) RNA-seq analysis of protein coding genes in mouse NPCs upon CRISPR/Cas9-mediatied disruption of Trim28. (d-e) PCA analysis of gene expression was unable to distinguish the Trim28-KO cells from Ctl, while PCA analysis of TE expression was.

**Supplementary figure 2.**
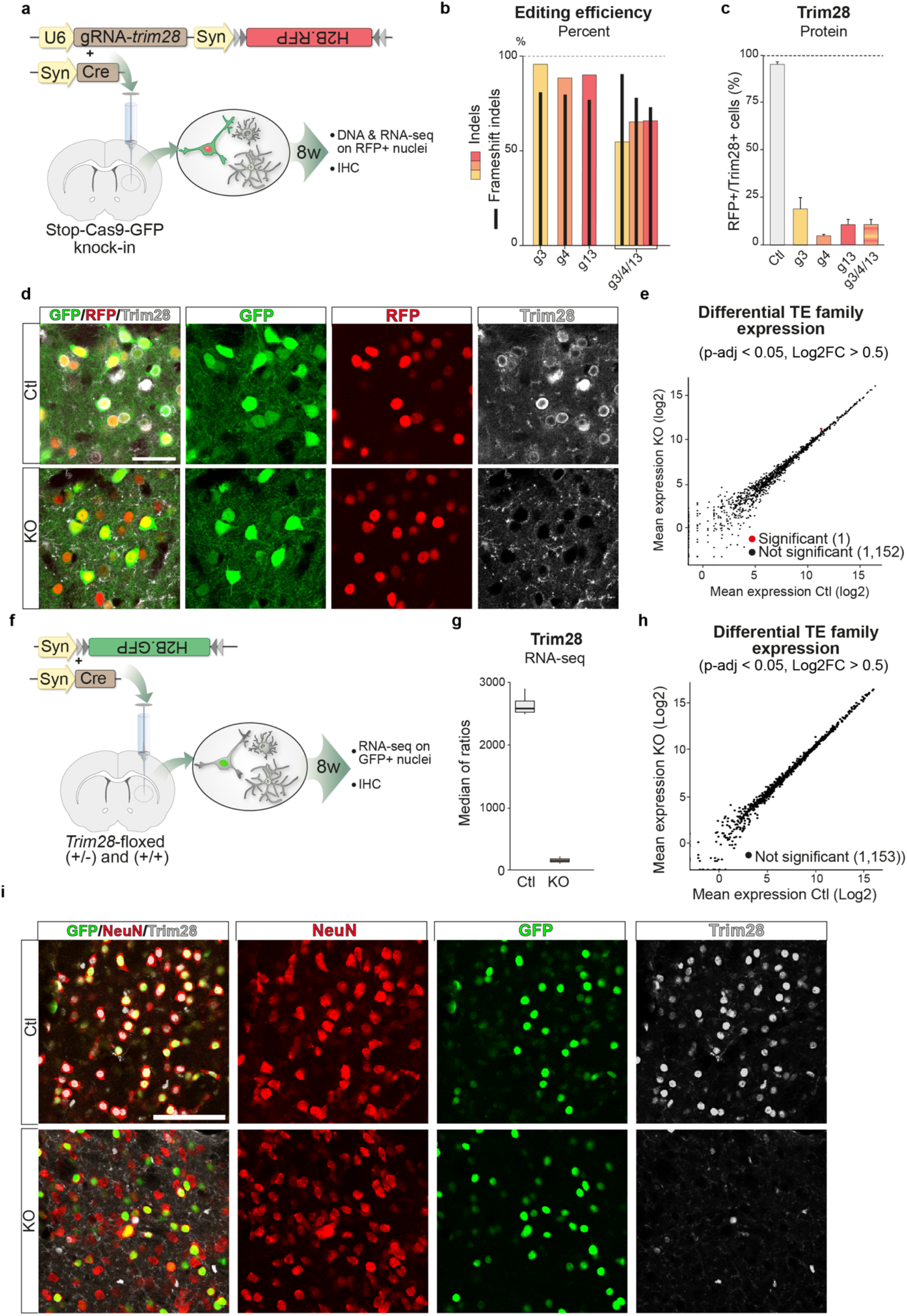
(a) A schematic of the workflow targeting Trim28 in the forebrain of Stop-Cas9-GFP knock-in mice using AAV vectors expressing the gRNA and a nuclear RFP reporter as well as an AAV vector expressing Cre, specifically in neurons by the Synapsin promoter. 8 weeks post injection, the animals were analysed either by IHC or by DNA/RNA sequencing following nuclei isolation by FACS. (b) Gene editing efficiency was evaluated by amplicon sequencing of the respective targeted sequences. Black bars indicate % of frameshift mutations. (c-d) Neuron specific editing of the Trim28-loci resulted in a robust loss of the Trim28 protein in neurons, as evaluated by IHC where the expression of Trim28 in RFP+ cells were quantified. Scalebar 30μm (e) RNA-seq analysis of TE expression using TEtranscripts in adult neurons from the Stop-Cas9-GFP knock-in mice upon Trim28-KO. (f) A schematic of the workflow targeting Trim28 in adult neurons in the forebrain of Trim28-flox mice (+/− and +/+) by co-injecting AAV vectors expressing Cre or nuclear GFP under the control of the Synapsin promoter. 8 weeks post injection, the animals were analysed either by IHC or by RNA sequencing following nuclei isolation by FACS. (g) RNA-seq analysis of the isolated GFP+ nuclei revealed a loss of the Trim28 transcript. (h) RNA-seq analysis of TE expression using TEtranscripts on the adult neurons in the Trim28-flox animals. (i) IHC for Trim28, GFP and the neuronal marker NeuN revealed a complete neuronal loss of the protein Trim28 in the Trim28-KO animals (Trim28-flox mice (+/+)). Scalebar 75μm.

**Supplementary figure 3.**
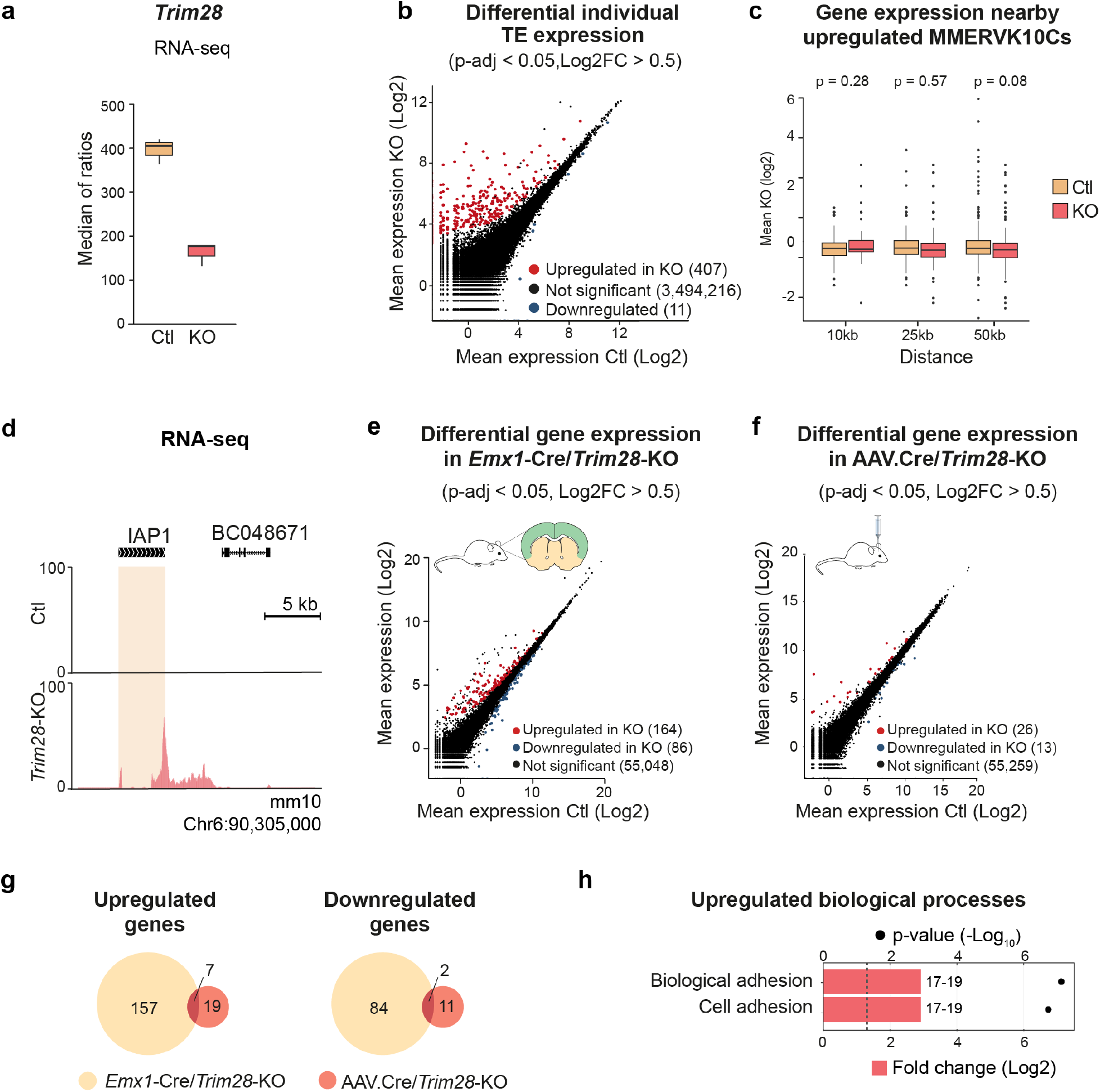
(a) RNA-seq analysis of Trim28 expression in the adult cortex of Emx1-Cre/ Trim28-KO animals (Emx1-Cre(+/−), Trim28-flox(+/+)). (b) RNA-seq analysis for iindividual TE elements in the adult cortex of Emx1-Cre/Trim28-KO animals compared to their Cre-negative control litter mates. (c) Expression of genes nearby upregulated full-length MMERVK10Cs were not upregulated in the Emx1-Cre/Trim28-KO animals. x-axis indicate the window-of-inclusion for genes located close to an MMERVK10C. (d) No transcriptional readthrough into nearby genes from upregulated MMERVK10Cs were observed, exemplified with an UCSC screenshot of the same location in the genome as shown in Fig 1f. (e) Differential gene expression in the adult cortex of Emx1-Cre/Trim28-KO animals compared to their control litter mates. (f) Differential gene expression in the adult forebrain upon AAV.Cre/Trim28-KO compared to controls. (g) Venn diagrams showing significantly up- and downregulated genes in the Emx1-Cre/Trim28-KO and the AAV.Cre/Trim28-KO animals, as well as the genes overlapping in between them. (h) GO term analysis of the overlapping upregulated genes between Emx1-Cre/Trim28-KO and the AAV.Cre/Trim28-KO animals.

**Supplementary figure 4.**
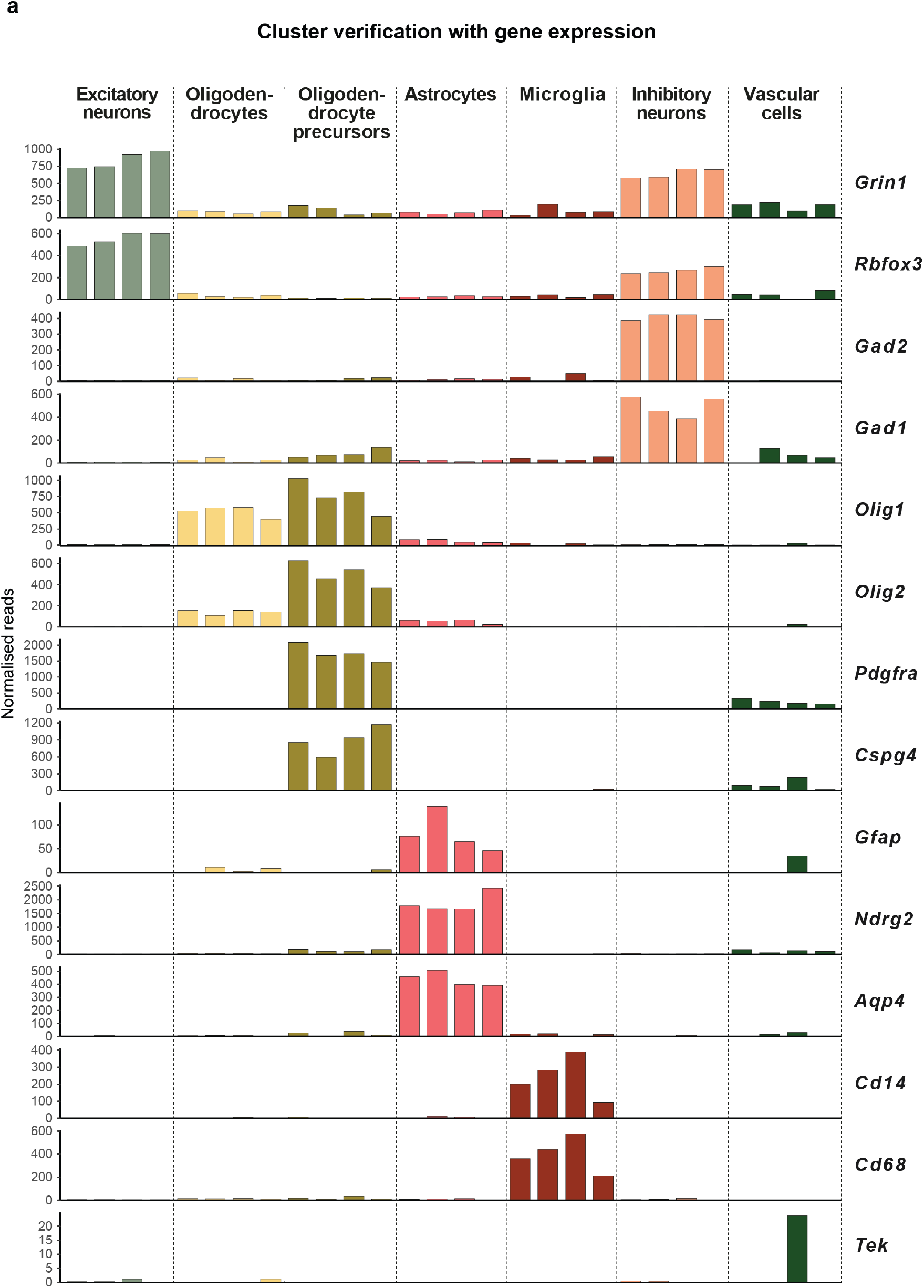
(a) The different cell types in the various clusters were verified by their expression of cell specific markers.

**Supplementary figure 5.**
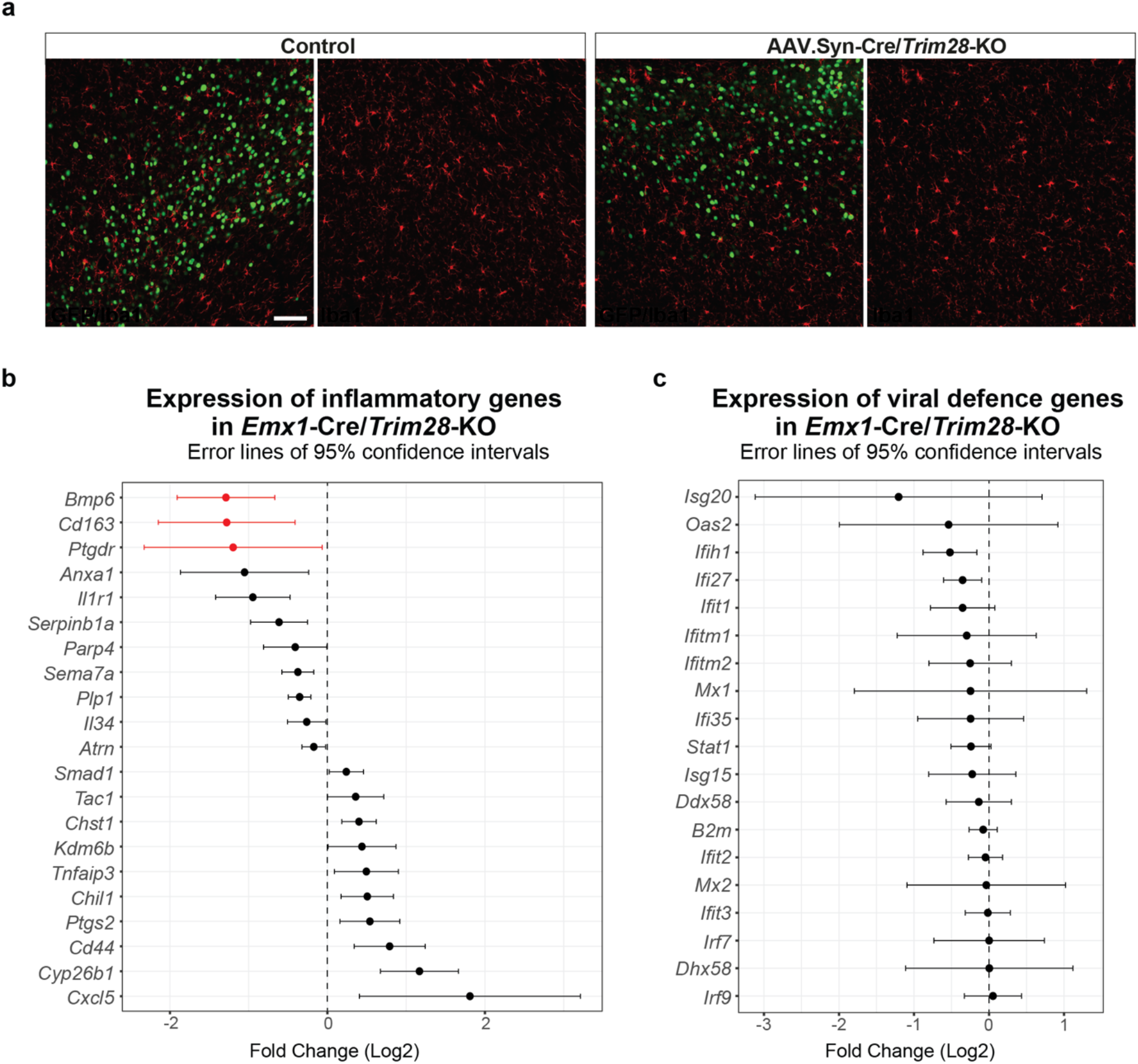
(a) Immunohistochemical analysis for the microglia marker upon Trim28 KO in mature neurons in vivo (AAV.Syn-Cre/Trim28-KO animals), revealed no difference in Iba1 morphology between Trim-28 KO animals and controls. Scalebar 75 μm. (b) Inflammatory genes were chosen from 216 genes annotated in the immune response GO term (GO0006954) if none of the confidence intervals overlapped zero Three genes were significantly downregulated upon the Emx1-Cre/Trim28-KO and are labelled in red: Bmp6, Cd163 and Ptgdr (p-adj < 0.05). c) The list of viral defence genes were taken from (Liu et al., 2018). None of these were significantly different upon the Emx1-Cre/Trim28-KO.

